# Distribution of the invasive *Anisandrus maiche* (Coleoptera: Scolytinae) in Switzerland, first record in Europe of its ambrosia fungus *Ambrosiella cleistominuta*, and its new association with *Xylosandrus crassiusculus*

**DOI:** 10.1101/2023.03.30.534995

**Authors:** José Pedro Ribeiro-Correia, Simone Prospero, Ludwig Beenken, Peter H. W. Biedermann, Simon Blaser, Yannick Chittaro, David Frey, Doris Hölling, Sezer Olivia Kaya, Miloš Knížek, Jana Mittelstrass, Manuela Branco, Beat Ruffner, Andreas Sanchez, Eckehard G. Brockerhoff

## Abstract

In 2022, two independent insect surveys in canton Ticino (southern Switzerland) revealed the widespread occurrence of the invasive ambrosia beetle *Anisandrus maiche* from southern to central-upper Ticino. This species is native to east Asia and has previously been found as a non-native invasive species in the United States, Canada, western Russia, Ukraine and, in 2021, in northern Italy. Here, we present the results of several trapping studies using different trap types (bottle traps, funnel traps and Polytrap intercept traps) and attractants and a map of the distribution of the species. In total, 685 specimens of *A. maiche*, all female, were trapped, and the identity of selected individuals was confirmed by morphological and molecular identification based on three mitochondrial and nuclear markers (COI, 28S and CAD). Traps checked from early April to early September 2022 in intervals of two to four weeks showed that flights of *A. maiche* occurred mainly from June to mid-August. Isolation of fungal associates of *A. maiche* from beetles trapped alive revealed the presence of four fungal species, including the ambrosia fungus *Ambrosiella cleistominuta*, the known mutualists of *A. maiche*. The identity of *A. cleistominuta* was confirmed by comparing DNA sequences of its nuclear, internal transcribed spacer (ITS) gene with reference sequences in NCBI and BOLDSYSTEMS. This represents the first record of *A. cleistominuta* in Europe. *Ambrosiella cleistominuta* was also found in association with another non-native invasive ambrosia beetle, *Xylosandrus crassiusculus*, at a botanic garden in central Ticino. As ambrosia beetles usually show a high degree of fidelity with only one mutualistic fungus (in the case of *X. crassiusculus* normally *Ambrosiella roeperi*), this association is highly unusual and probably the result of lateral transfer among these non-native invasive species. Of the other fungal associates isolated from *A. maiche* in Ticino, *Fusarium lateritium* is of note as there is a possibility that *A. maiche* could act as a vector of this plant pathogen. We highlight several research needs that should be addressed to gain insight into the potential impact of these non-native species and to overcome problems with heteroplasmy in COI sequences in studies of invasion and population genetics of ambrosia beetles.

## Introduction

Biological invasions are a growing concern due to the continuing increase in establishments of non-native (alien) invasive species and their impacts on native species, natural and modified ecosystems and on plant, animal and human health (Kenis et al. 2009; Brockerhoff et al. 2017; Seebens et al. 2018; Pyšek et al. 2020). Bark and ambrosia beetles (Coleoptera: Curculionidae, Scolytinae) are particularly successful as invaders because they are often transported internationally by trade in wood products and because of the widespread use of wood packaging materials such as pallets and crates (Brockerhoff et al. 2006; Lantschner et al. 2020). In addition, many ambrosia beetles have mating systems that involve inbreeding (Kirkendall 1983) which greatly facilitates the establishment of small invading populations (Kirkendall and Faccoli 2010; Lantschner et al. 2020). Furthermore, because ambrosia beetles typically feed on specific ambrosia fungi which they carry in their mycangia and cultivate in galleries they construct in deadwood, these beetles often have a very wide host range (e.g., Ranger et al. 2016; Hulcr and Stelinski 2017). This also contributes to their invasiveness since they are less dependent on the presence of a particular host species. Although native ambrosia fungi are typically not pathogenic, some non-native ambrosia beetles are vectors of severe plant pathogens that can cause tree death. For example, the Asian *Xyleborus glabratus* Eichhoff, 1877 and its symbiotic tree pathogen *Raffaelea lauricola* T.C. Harr., Fraedrich & Aghayeva (Harrington et al. 2008), the causal agents of laurel wilt, cause large-scale mortality of many trees in the family Lauraceae in the southeastern United States (Fraedrich et al. 2008; Hughes et al. 2017).

*Anisandrus maiche* (Kurentzov, 1941) is an invasive ambrosia beetle native to northeast Asia (parts of China, Japan, North and South Korea, and the Russian Far East (Knížek 2011; Terekhova & Skrylnik 2012; Alonso-Zarazaga et al. 2017, 2023; Smith et al. 2020; EPPO 2022). Established non-native populations of *A. maiche* were detected in North America first in 2005 in Pennsylvania, USA, and subsequently in the US states of Ohio, West Virginia, Illinois, Indiana, Maryland, New Jersey, New York, and Wisconsin, and in the Canadian Provinces Ontario and Quebec (Rabaglia et al. 2009; Haack et al. 2013; Gomez et al. 2018; Thurston et al. 2022). In Europe, non-native populations were detected first in 2007 in western Russia and in eastern Ukraine (Moscow oblast, Belgorod oblast, Donetsk oblast, Kharkiv oblast and Sumy oblast (Nikulina et al. 2007; Nikitskii 2009; Terekhova & Skrylnik 2012; Kovalenko & Nikitski 2013; Nikulina et al. 2015)). About 14 years later, in 2021-2022, this beetle was found in Italy’s Veneto and Lombardy regions (Colombari et al. 2022; Ruzzier et al. 2022). In July 2022, several specimens of *A. maiche* were identified in trap catches from near Biasca, Canton Ticino, in southern Switzerland. Subsequently, many more specimens were found in trap catches from several other locations across the southern half of Canton Ticino and near Roveredo in southwestern Canton Grisons, suggesting that the species has been established in these areas for several years.

Here, our objectives are to report on the discovery of *Anisandrus maiche* in Switzerland, the locations and forest types where it was found, and the traps and attractants with which the species was caught. Furthermore, we provide information on the fungal and microbial associates which we recorded from *A. maiche* in Switzerland, including its ambrosia fungus *Ambrosiella cleistominuta* C. Mayers & T.C. Harr, and on potential damage caused by this beetle, based on a review of available information. In addition, we report an unusual association of *Xylosandrus crassiusculus* (Motschulsky 1866) with the ambrosia fungus of *Anisandrus maiche*.

## Methods and materials

### *Study sites of* A. maiche *in cantons Ticino and Grisons, and description of traps and attractants*

All study sites in Switzerland where *A. maiche* was found are located in the southern part of the country in the cantons Ticino (40 sites) and Grisons (one site, Roveredo) (**Table 1, Fig. 1**). The sites were in mixed forests composed mostly of sweet chestnut (*Castanea sativa*), oaks (*Quercus* sp.), Scots pine (*Pinus sylvestris*) and other trees, with varying proportions of broadleaved trees and conifers.

**Table 1:**
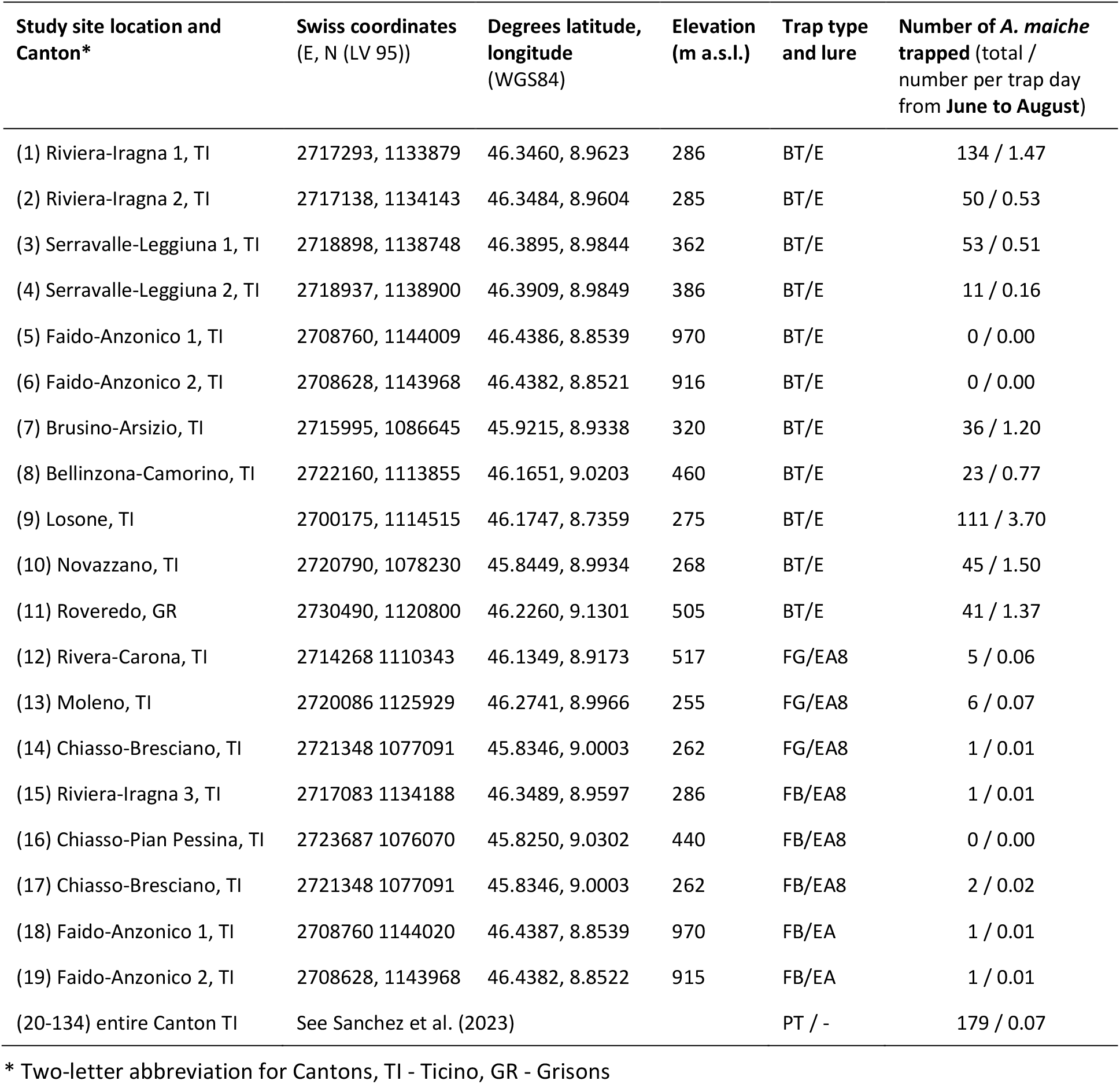
Study site locations, traps used and the number of *Anisandrus maiche* trapped in 2022. Trap types: BT, bottle trap; FG, funnel trap green; FB, funnel trap black; PT, Polytrap. Lures: E, ethanol; EA, ethanol + alpha-pinene; EA8, ethanol + alpha-pinene + eight-component blend (for details about trap types and lures see methods).

**Figure 1.**
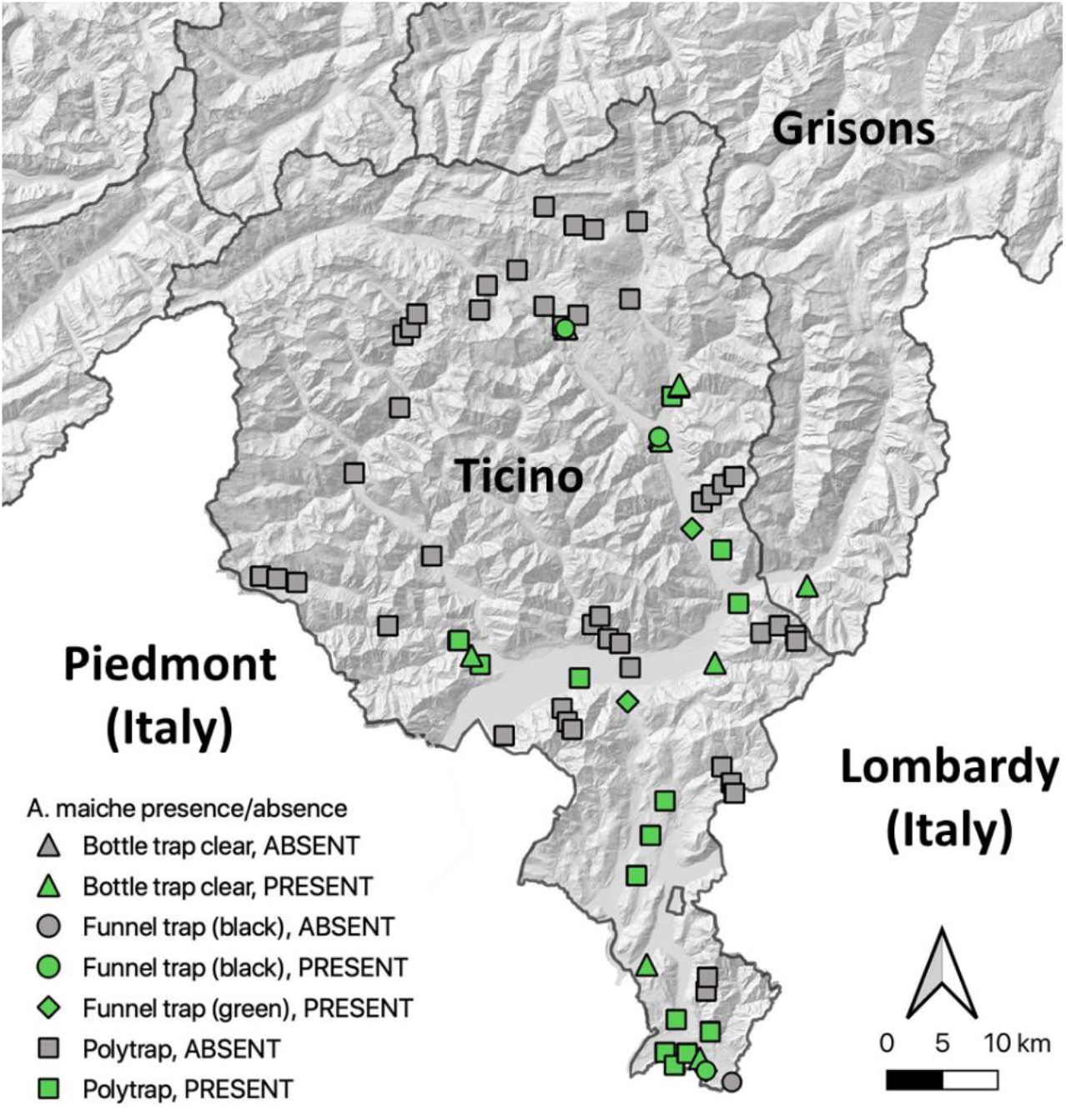
Trap locations in cantons Ticino and Grisons where *Anisandrus maiche* was captured (green symbols) or where no captures were recorded (grey symbols). Symbols vary by trap type (see legend and methods for details). Note that each square represents a pair of two Polytrap intercept traps which were placed in close proximity to each other (Vector and raster map data © swisstopo.ch).

Five trap type and lure combinations were used in 2022 during several insect surveys where *A. maiche* was caught. **Type 1** (bottle traps, “**BT**”) traps consisted of bottle traps made at the laboratory based on the design described in Grégoire et al. (2003) and as modified by Gossner et al. (2019) with 70% ethanol as lure and propylene glycol as preservative and installed such that the bottle part was about 1 m above ground. Two BT were installed at each study site (see Table 1) on 2 April 2022 and taken down on 26 August 2022. A small number of additional BT traps were used without propylene glycol to enable live captures for studies of fungal associates (see below).

**Type 2** (funnel traps green, “**FG**”) traps consisted of green multi-funnel traps (ChemTica Internacional, Costa-Rica) with propylene glycol as preservative and baited with alpha-pinene and ethanol (Econex, Spain) and an 8-component lure blend (fuscumol, fuscumol acetate, geranylacetone, monochamol, 3-hydroxyhexan-2-one, anti-2,3-hexanediol, 2-methylbutan-1-ol, and prionic acid) as described by Fan et al. (2019). FG traps were suspended by a rope inside the canopy of trees at a height of approximately 10 m. One of these traps was installed at each study site (see Table 1) on 12 April **2021** and taken down on 1 or 29 September **2021**.

**Type 3** (funnel traps, black, “**FB**”) traps consisted of black multi-funnel traps with propylene glycol as preservative and the same lure combination and trapping period as Type 2. FB traps were suspended by a rope below the canopy of trees at a height of 2-3 m above ground. One of these traps was installed at each study site (see Table 1) on 12 April **2021** and taken down on 1 or 29 September **2021**. Trapping with funnel traps (FG and FB) in 2021 was undertaken as part of a surveillance programme aimed at priority quarantine pests targeting mainly longhorn beetles.

**Type 4** (Polytrap, “**PT**”) traps were unbaited Polytrap™ interception traps (École d’Ingénieurs de Purpan, Toulouse, France; Brustel 2012) with saturated salt solution and neutral detergent as preservative, suspended 2 m above the ground. As part of a biodiversity survey funded by the Canton Ticino (MSNL and UPN), a total of 114 Polytraps were installed on 10 March 2022 and taken down at the end of 3 October 2022 (see Sanchez et al. (2023) for details).

In addition to the traps described above, collection of ambrosia beetles was also attempted with log sections of European beech (*Fagus sylvatica*), Norway spruce (*Picea abies*) and sweet chestnut ca. 50 cm long, 5-10 cm diameter, baited with 70% ethanol and suspended alongside tree stems at a height of 1.5-2.0 m, as described by Monterrosa et al. (2021). These were installed at study sites (1)-(6) in Ticino (Table 1) from 16 April to 26 August 2022. In addition, short sections of beech branches (about 20 cm long, 2-4 cm diameter) soaked in 70% ethanol were placed twice on the ground at study sites (1)-(6) in Ticino from 12 June to 25 July 2022.

### Study sites in cantons Valais and Zurich

Apart from traps placed in cantons Ticino and Grisons, bottle traps with ethanol as lure (as trap type 1 described above) were also used in canton Valais from 18 March 2022 – 25 August 2022 at six locations (Brig 1, 46.2905°N, 7.9601°E, 1260 m a.s.l.; Brig 2, 46.2941°N, 7.9577°E, 1259 m a.s.l.; Lens 1, 46.2677°N, 7.4339°E, 1096 m a.s.l.; Lens 2, 46.2677°N, 7.4339°E, 1097 m a.s.l.; Visp 1, 46.2971°N, 7.8566°E, 676 m a.s.l.; and Visp 2, 46.2967°N, 7.8564°E, 705 m a.s.l.), and in canton Zurich from 4 April 2022 – 31 August 2022 at six locations (Zurich-Hönggerberg 1, 47.4121°N, 8.4978°E, 535 m a.s.l.; Zurich Hönggerberg 2, 47.4196°N, 8.4872°E, 525 m a.s.l.; Stallikon-Uetliberg 1, 47.3364°N, 8.4936°E, 660 m a.s.l.; Stallikon-Uetliberg 2, 47.3367°N, 8.4945°E, 669 m a.s.l.; Birmensdorf-Rameren 1, 47.363°N, 8.4483°E, 540 m a.s.l.; Birmensdorf Rameren 2, 47.3631°N, 8.4483°E, 555 m a.s.l.). The study sites in Cantons Valais and Zurich were also in mixed forests with varying proportions of broadleaved trees and conifers and composed mostly of oaks, beech, Norway spruce, Scots pine or other trees.

### *Study site of* Xylosandrus crassiusculus *in Ticino*

Specimens of the granulate ambrosia beetle *X. crassiusculus* along with their breeding galleries and associated fungi were obtained from the Botanical Garden on the Brissago Island (46.13229°N, 8.73535°E, 205 m a.s.l.) following a report of unusual frass appearing on branches (diameter: 0.9 - 3 cm) of a *Hakea* sp. (Proteaceae) shrub. Several adults and larvae of *X. crassiusculus* were isolated from the *Hakea* material on 1 September 2022.

### *Curation and identification of* A. maiche

All ambrosia and bark beetles from traps with propylene glycol preservative were sorted under a stereomicroscope and kept in 70% ethanol for temporary storage while selected individuals were pinned. Ambrosia and bark beetles were identified morphologically by J. Ribeiro-Correia, E. Brockerhoff, A. Sanchez and M. Knížek using Grüne (1979), Pfeffer (1995), Stark (1952), Rabaglia et al. (2009) and reference collections held at WSL, by M. Knížek and by Andreas Sanchez. A specimen of *A. maiche* from Serravalle-Leggiuna 1 is shown in Fig. 2.

**Figure 2.**
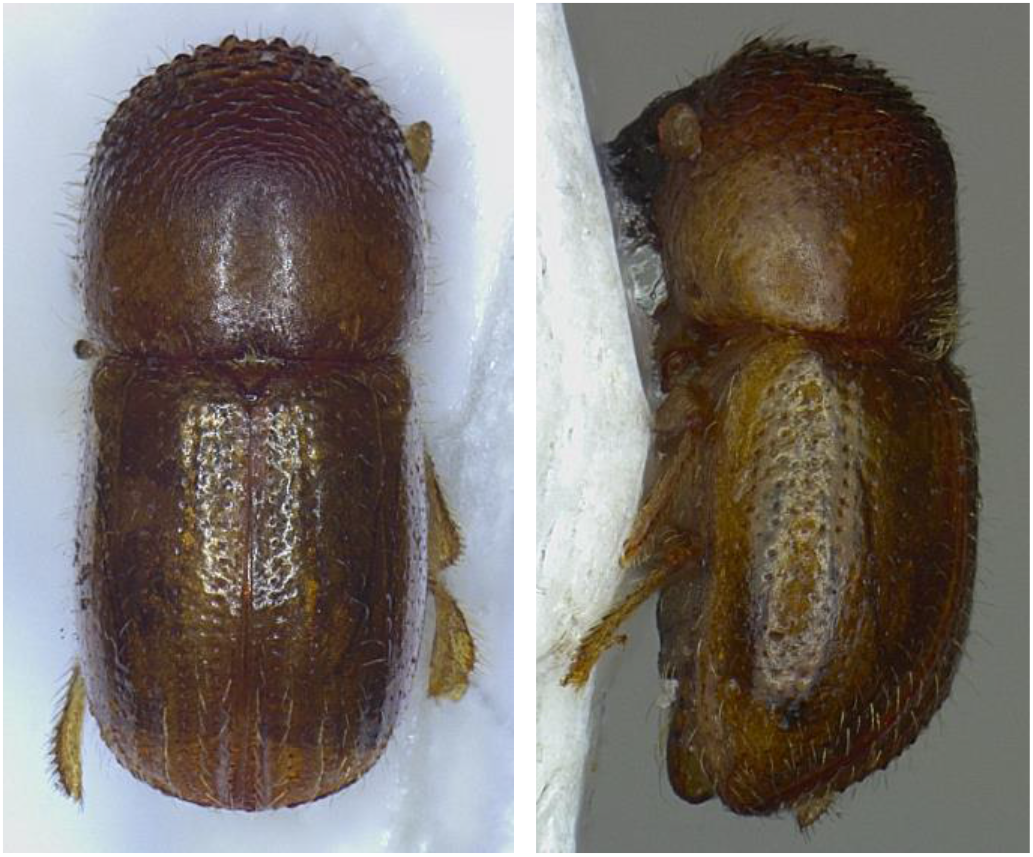
*Anisandrus maiche* adult trapped at Serravalle-Leggiuna 1 (Ticino), dorsal and lateral. Photos by Carl-Michael Anderson, WSL. (L1_13.5.22_AM_sm_1 / L1_13.5.22_AM_sm_1_lateral).

For molecular diagnostics, genomic DNA from *A. maiche* was extracted from adults using the NucleoSpin® Tissue XS Kit (Macherey-Nagel, Düren, Germany) using whole insects or, to preserve the specimens, leg fragments and one elytron. The COI barcode region was amplified and sequenced with primers LCO1490/HCO2198 (Folmer et al., 1994). The nuclear markers, the ribosomal-encoding gene 28S and the CAD gene were targeted with primers 3690s/a4285 (Kambestad et al. 2017) and CADforB2/apCADrevlmod (Dole et al. 2010), respectively. Sequences were trimmed and assembled using the CLC Main Workbench Version 22.0.2 (QIAGEN, Aarhus, Denmark) and checked manually before being subjected to BLAST searches against the All Barcode database on BOLD (http://www.boldsystems.org) and NCBI (https://blast.ncbi.nlm.nih.gov). All COI barcode sequences from *A. maiche* generated in this study are deposited on BOLD (accession numbers listed below). In addition, a specimen of *A. maiche* (BOLD Sequence ID: SCOL295-12) collected in Primorsky Krai, Russian Far East, was obtained from Bjarte Jordal (University of Bergen, Natural History Collections) and processed for molecular analyses as described above for comparison with our specimens.

Voucher specimens are held at WSL, Birmensdorf, Switzerland, at the Museum of Natural History, Lugano, Switzerland, and in the collections of M. Knížek in Prague, Czechia, A. Sanchez in Sion, Switzerland, and Heiko Gebhardt in Tübingen, Germany. Details on individual reference specimens are provided below.

### *Extraction, cultivation, and identification of fungal species associated with* A. maiche

Live bottle traps (as described above for trap type 1, but with moistened paper instead of propylene glycol in the collection jar) were installed at six locations in Ticino (locations number (1)-(6), see Table 1) to collect live beetles for extraction and cultivation of associated fungi. Traps were inspected after 2 to 4 days in the field. Beetles present in the traps were removed immediately from the PVC bottle, placed individually in 1.5 mL Eppendorf tubes and, upon arrival at the laboratory, stored at 4°C for a maximum of one week.

The fungal species present on the surface and inside the collected beetles were identified as follows. First, the individual beetles were extracted from the Eppendorf tubes using sterile tweezers and gently placed onto weaker-strength agar medium (SMA; 10g/L malt extract; 15g/L agar; 100 ppm streptomycin added after autoclaving to prevent growth of bacteria). The beetles were allowed to walk freely for 30-45 min so that fungal spores present on their body would eventually deposit on the agar surface. To identify fungal species inside their body (i.e., in the mycangia and in the digestive tract), beetles were subsequently removed from the Petri dish using sterile tweezers, placed in 90% ethanol for 1-2 seconds to kill any spores still present externally on their body, rinsed twice in distilled sterile water, and put on a sterile paper towel to dry. Once dry, beetles were placed individually in a new 1.5 mL Eppendorf tube containing 0.5 mL distilled sterile water and crushed with a sterile rod. After brief vortexing, 100 µL of this solution was spread on SMA and incubated in the dark at room temperature. Plates were checked daily for up to one week and growing fungal colonies were subcultured on Potato Dextrose Agar (PDA; 39 g l^-1^, Difco, Voight Global Distribution, Lawrence, MD, USA). When morphologically different colonies were present on a plate, a representative colony of each morphotype was transferred to PDA. After incubation of the PDA plates for two weeks in the dark at room temperature, fungal cultures were grouped into morphotypes based on the macro-morphological features of their mycelia.

For species identification, DNA was extracted from 1-3 representative cultures of each morphotype using LGC reagents and Kingfisher 96/Flex (LGC Genomics GmbH, Berlin, Germany), according to the manufacturer’s instructions. The nuclear, internal transcribed spacer (ITS) was then amplified by PCR and sequenced in both directions using the forward ITS5 and reverse ITS4 primers (White et al. 1990) using the general methodology described in Franić et al. (2019). Sequences were assembled and edited using CLC Main Workbench Version 22.0.2 and compared with reference sequences in NCBI and BOLDSYSTEMS databases. Two sequences were considered to belong to the same species if they showed at least 99% similarity.

### *Curation and identification of* X. crassiusculus *and extraction, cultivation, and identification of associated fungal species*

*Xylosandrus crassiusculus* (Fig. 3) obtained from the Brissago Island were identified morphologically by S. Blaser and by using the same molecular diagnostics protocol described above. For the isolation of the associated fungal species, infested branches (Fig. 3) were split lengthwise. From the breeding galleries, mycelial threads were isolated from the walls using a sterile needle and plated on Petri dishes containing DiaMalt agar with streptomycin (15 g Plant Propagation Agar (Conda), 20 g DiaMalt (Hefe Schweiz AG), 100 mg streptomycin (Sigma), 1 l ddH2O).

**Figure 3.**
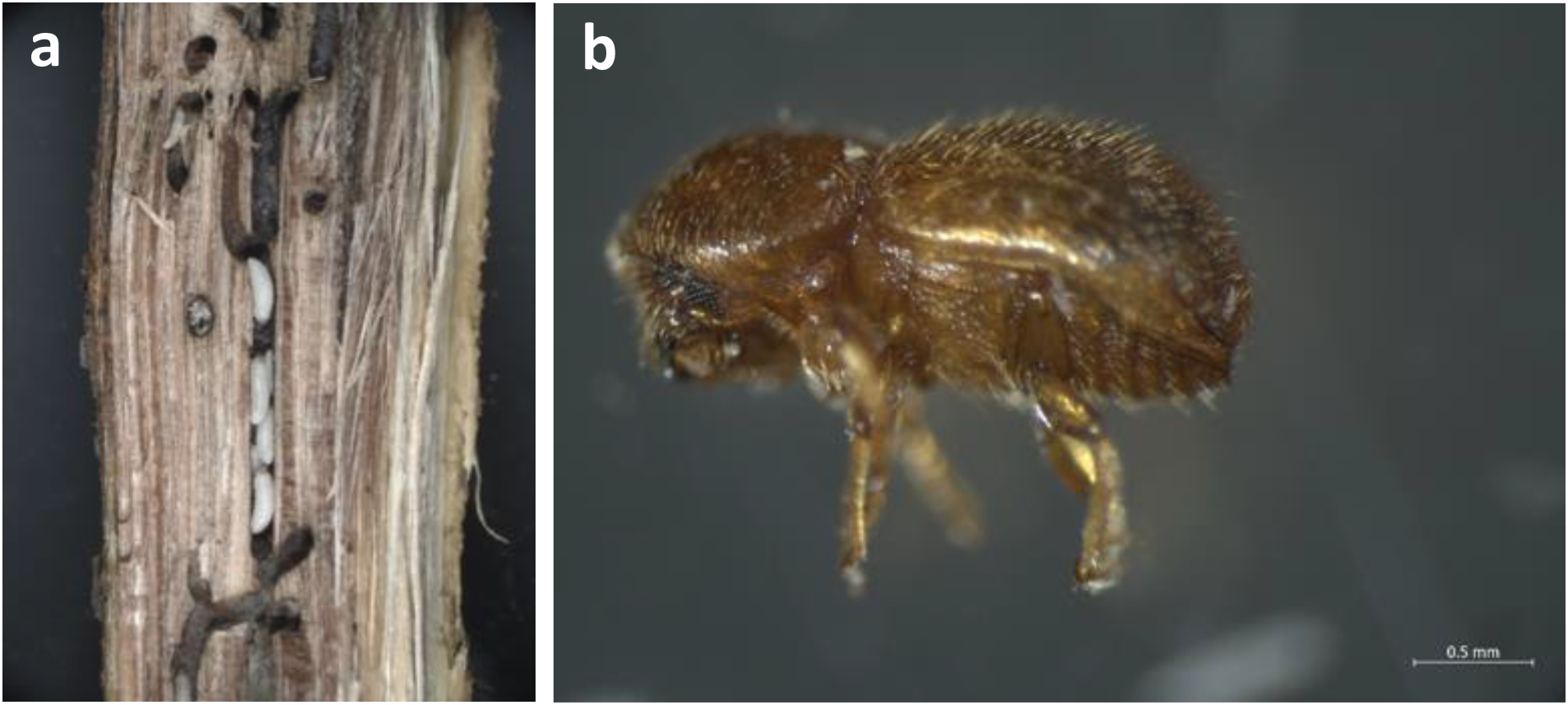
*Xylosandrus crassiusculus* larvae in galleries in a *Hakea* sp. stem (a) and a young adult beetle (b). Photos by Simon Blaser, WSL

The fungus was also isolated from one living beetle by grasping the beetle with sterile tweezers and pressing the pronotum several times into a Petri dish with water agar with streptomycin (15 g Plant Propagation Agar (Conda), 100 mg streptomycin (Sigma), 1 l ddH2O). The pronotum was pressed into the agar in such a way that the mycangium opened. The growing fungal colonies were sub-cultured on DiaMalt agar with streptomycin as soon as they were visible after some days. Fungal DNA was isolated from the mycelia and amplified and sequenced using the method described above.

## Results

### *Specimens of* A. maiche *trapped in Switzerland*

In 2022, a total of 685 specimens of *A. maiche* were trapped in southern Switzerland (Table 1, Fig. 1), including 504 in bottle trap samples and 179 in Polytrap samples. The first specimens were discovered in bottle trap catches from May 2022 from the vicinity of Biasca (Riviera-Iragna and Serravalle-Leggiuna), which were identified by Miloš Knížek on 1 July 2022 and confirmed by molecular analysis soon after (see below). Over the summer of 2022, 248 specimens of *A. maiche* were trapped with bottle traps (all with ethanol as lure) at Riviera-Iragna and Serravalle-Leggiuna (Table 1). An additional 256 *A. maiche* were trapped at five locations in southern Ticino with additional traps placed there to determine the extent of the distribution of *A. maiche* in Ticino (Table 1).

Two more specimens of *A. maiche* were found in samples from a black funnel trap from Faido-Anzonico in northern Ticino, the northernmost occurrence of the species (Table 1). This was part of another study which used funnel traps similar to trap type 3 (FB) but baited only with ethanol and alpha-pinene lures and suspended from pine branches about 2-3 m above ground.

Polytraps (trap type 4) installed in 2022 across canton of Ticino caught 179 specimens of *A. maiche*. These were captured with 25 traps distributed between Chiasso (southernmost Ticino) and Biasca (central-upper Ticino) out of a total of 114 polytraps placed across Ticino. No specimens were captured in the Polytraps located north of Biasca and in the Maggia Valley north of Terre di Pedemonte (near Ascona).

Following the discovery of *A. maiche* in the vicinity of Biasca in June-July 2022, samples collected in a Swiss surveillance programme in 2021 (using green and black funnel traps, trap types 2 and 3) aimed at detecting priority quarantine insects, especially longhorn beetles (Cerambycidae), were re-examined for the presence of *A. maiche*. In these samples, a total of 15 *A. maiche* were found in southern Ticino near the Italian border (two sites near Chiasso), north of Lugano (Rivera), north of Bellinzona (Moleno), and near Biasca (Riviera-Iragna) (Table 1, Fig. 1).

Across all sites and trap types, most specimens of *A. maiche* were trapped at lower elevations in the valleys or lower mountain slopes at elevations between 195 m a.s.l. (near Locarno) and 386 m a.s.l. (near Biasca), but a few were caught at higher elevations such 626 m a.s.l. (Capriasca, north of Lugano) and 916 m a.s.l. (Faido Anzonico). All specimens were trapped in a variety of forest types with sweet chestnut, beech, mixed broadleaved trees and, in a few cases, a mixture with Scots pine.

Bottle trap captures of ambrosia beetles from six sites in canton Valais and six sites in canton Zurich (that were part of the same study described here) did not reveal the presence of *A. maiche*.

### *Molecular identification of* A. maiche *and* X. crassiusculus

To confirm the identity of selected specimens of *A. maiche*, nucleotide BLAST searches were performed on BOLD and NCBI (as accessed in October 2022). For the mitochondrial COI barcode region, seven out of our 14 samples share 100% identity with accession MN619845 on NCBI, designated as *A. maiche*. For three other samples from Switzerland, a second haplotype was identified sharing 100% identity with a private accession on BOLD, also designated as *A. maiche*. The two haplotypes show a remarkable number of base substitutions between haplyotypes generating a divergence of 5.9%. In addition, four specimens show a pattern of heteroplasmy, compatible with the two haplotypes identified. However, assessing the nuclear, ribosomal-encoding gene 28S revealed 100% identity for all specimens to the ribosomal-encoding gene 28S of *A. maiche* (GenBank Accession MK098863, voucher specimen UFIFAS UFFE 28176). In addition, the CAD fragment from three samples (PHP22_0410, PH22_0411, PHP22_0539) displayed 100% identity to a sequence of *A. maiche* (GenBank Accession MN260139, Shanghai, China), but differed by 1bp to accession MN260138 collected in Michigan, USA. The *A. maiche* specimen from the Russian Far East (BOLD Sequence ID: SCOL295-12) which was re-analysed together with our specimens from Switzerland showed the same pattern of COI heteroplasmy and shared 100% identity to the 28S locus of all our analysed specimens.

DNA barcoding analyses to confirm the identity of the ambrosia beetle sample obtained from the gallery in a *Hakea* shrub from the botanical garden on Brissago Island clearly showed that this was *X. crassiusculus* (WSL DNA-ID PHP22_0813). In a BOLD blast analysis, the specimen shared an identity of 99.8% with a reference sequence of *X. crassiusculus* (NCBI GenBank accession number MN620076).

### List of selected specimens held in collections

Selected specimens from Switzerland, collected by José Ribeiro-Correia, caught in bottle traps with ethanol as lure, along with GenBank accession numbers:

2 females, Leggiuna 1 (Serravalle-Leggiuna), Ticino, LV95: 2718898 E, 1138748 N (46.3895°N, 8.9844°E), 370 m a.s.l., 13 - 27 May 2022, det. Miloš Knížek, WSL DNA-IDs PHP22_0410 (GenBank: OQ685554) and PHP22_0411 (GenBank: OQ685554), entire specimens used for molecular analysis, DNA held at WSL PHP.

1 female, Iragna 1 (Riviera-Iragna), Ticino, LV95: 2717293 E, 1133879 N (46.3460°N, 8.9623°E), 286 m a.s.l., 27 May – 10 June 2022, det. Miloš Knížek, WSL DNA-ID PHP22_0539 (GenBank: OQ685552), entire specimens used for molecular analysis, DNA held at WSL PHP.

7 females, Iragna 2 (Riviera-Iragna), Ticino, LV95: 2717138 E, 1134143 N (46.3484°N, 8.9604°E), 285 m a.s.l., 10 – 25 June 2022, det. José Correia, WSL DNA-IDs PHP22_0627 (GenBank: OQ685551), PHP22_0628 (GenBank: OQ685550), PHP22_0629 (GenBank: OQ685549), PHP22_0630 (GenBank: OQ685548), PHP22_0946 (GenBank: OQ685547), PHP22_0947(GenBank: OQ685546), PHP22_0952(GenBank: OQ685541), specimens at WSL FPS (ethanol/freezer), DNA held at WSL PHP.

1 female, Brusino-Arsizio, Ticino, LV95: 2715995 E, 1086645 N (45.9215°N, 8.9338°E), 320 m a.s.l., 18 July – 3 Aug. 2022, det. José Correia, WSL DNA-ID PHP22_0948 (GenBank: OQ685545), specimen at WSL FPS (ethanol/freezer), DNA held at WSL PHP.

3 females, Roveredo, Grisons, LV95: 2730490 E, 1120800 N, (46.2260°N, 9.1301°E), 505 m a.s.l., 18 July – Aug. 2022, det. José Correia, WSL DNA-IDs PHP22_0949 (GenBank: OQ685544), PHP22_0950 (GenBank: OQ685543), PHP22_0951 (GenBank: OQ685542), specimens at WSL FPS (ethanol/freezer), DNA held at WSL PHP.

2 females, Leggiuna 1 (Serravalle-Leggiuna), Ticino, LV95: 2718898 E, 1138748 N (46.3895°N, 8.9844°E), 370 m a.s.l., 20 April – 13 May 2022 / 13-27 May 2022, det. Miloš Knížek, specimens at WSL FPS (pinned).

3 females, Leggiuna 1 (Serravalle-Leggiuna), Ticino, LV95: 2718898 E, 1138748 N (46.3895°N, 8.9844°E), 370 m a.s.l., 13 – 27 May 2022, det. José Correia, specimens at WSL FPS (pinned).

2 females, Iragna 2 (Riviera-Iragna), Ticino, LV95: 2717138 E, 1134143 N (46.3484°N, 8.9604°E), 285 m a.s.l., 13 – 27 May 2022, det. Miloš Knížek, specimens at Miloš Knížek collection, Prague, Czechia.

11 females, Iragna 1 (Riviera-Iragna), Ticino, LV95: 2717293 E, 1133879 N (46.3460°N, 8.9623°E), 286 m a.s.l., 27 May – 10 June 2022, det. Miloš Knížek, specimens at Miloš Knížek collection, Prague, Czechia.

2 females, Leggiuna 1 (Serravalle-Leggiuna), Ticino, LV95: 2718898 E, 1138748 N (46.3895°N, 8.9844°E), 370 m a.s.l., 13 – 27 May 2022, det. Miloš Knížek, specimens at Miloš Knížek collection, Prague, Czechia.

1 female, Iragna 2 (Riviera-Iragna), Ticino, LV95: 2717138 E, 1134143 N (46.3484°N, 8.9604°E), 285 m a.s.l., 10 – 25 June 2022, det. Miloš Knížek, specimen at Miloš Knížek collection, Prague, Czechia.

1 female, Novazzano, Ticino, LV95: 2720790 E, 1078230 N (45.8449°N, 8.9934°E), 268 m a.s.l., 18 July – 3 Aug. 2022, det. Miloš Knížek, specimen at Miloš Knížek collection, Prague, Czechia.

1 female, Losone, Ticino, LV95: 2700175 E, 1114515 N (46.1747°N, 8.7359°E), 275 m a.s.l., 18 July – 3 Aug. 2022, det. Miloš Knížek, specimen at Miloš Knížek collection, Prague, Czechia.

Selected specimens from Switzerland, collected by David Frey, caught with unbaited Polytrap interception traps:

13 females, Ruderi del Castello di Claro (Bellinzona), Ticino, LV95: 2722751 E, 1124011 N (46.2563°N, 9.0306°E), 437 m a.s.l., 14 – 29 June 2022, det. Andreas Sanchez, specimens at MSNL.

7 females, Ruderi del Castello di Claro (Bellinzona), Ticino, LV95: 2722750 E, 1124033 N (46.2565°N, 9.0306°E), 453 m a.s.l., 3 -15 July –2022, det. Andreas Sanchez, specimens at MSNL.

1 female, Solorónch (Capriasca), Ticino, LV95: 2717651 E, 1101409 N (46.0539°N, 8.9588°E), 626 m a.s.l., 1 – 13 June 2022, det. Andreas Sanchez, specimen at MSNL.

2 females, Solorónch (Capriasca), Ticino, LV95: 2717643 E, 1101393 N (46.0538°N, 8.9587°E), 620 m a.s.l., 11 July – 15 August 2022, det. Andreas Sanchez, specimens at MSNL.

54 females, Bolette (Locarno), Ticino, LV95: 2709912 E, 1112488 N (46.1549°N, 8.8614°E), 195 m a.s.l., 17 May – 13 August 2022, det. Andreas Sanchez, specimens at MSNL and at Andreas Sanchez collection, Sion, Switzerland.

31 females, Bolette (Locarno), Ticino, LV95: 2709941 E, 1112476 N (46.1547°N, 8.8618°E), 195 m a.s.l., 2 June – 13 August 2022, det. Andreas Sanchez, specimens at MSNL and at Andreas Sanchez collection, Sion, Switzerland.

3 females, Colombera (Stabio), Ticino, LV95: 2717680 E, 1078726 N (45.8499°N, 8.9535°E), 339 m a.s.l., 1 – 27 June 2022, det. Andreas Sanchez, specimens at MSNL.

4 females, Ronco del Re (Terre di Pedemonte), Ticino, LV95: 2699033 E, 1115853 N (46.1868°N, 8.7213°E), 372 m a.s.l., 17 May – 8 August 2022, det. Andreas Sanchez, specimens at MSNL.

9 females, S. Martino (Vezia), Ticino, LV95: 2716328 E, 1098298 N (46.0262°N, 8.9409°E), 431 m a.s.l., 13 June – 12 August 2022, det. Andreas Sanchez, specimens at MSNL and at Heiko Gebhardt collection, Tübingen, Germany.

### Comparison of trap effectiveness

It was not an objective of the surveys reported here to compare the effectiveness of different trap types in capturing *A. maiche*. Nevertheless, a comparison of *A. maiche* captures per trap per day using bottle traps with ethanol as lure indicated that these were more effective than green or black funnel traps with ethanol and additional attractants (Table 1). Captures of *A. maiche* per trap per day during the main flight period (from June to August, see Fig. 4) at sites where the species occurred averaged 1.02 specimens per trap per day (n=11) for bottle traps with ethanol as lure (BT/E, trap type 1); 0.05 specimens per trap per day (n=3) for green funnel traps with ethanol, alpha-pinene and eight-component blend (FG/EA8, trap type 2); 0.01 specimens per trap per day (n=3) for black funnel traps with ethanol, alpha-pinene and eight-component blend (FB/EA8, trap type 3); and 0.07 specimens per trap per day (n=28) for Polytrap with no lure (PT, trap type 4).

**Figure 4.**
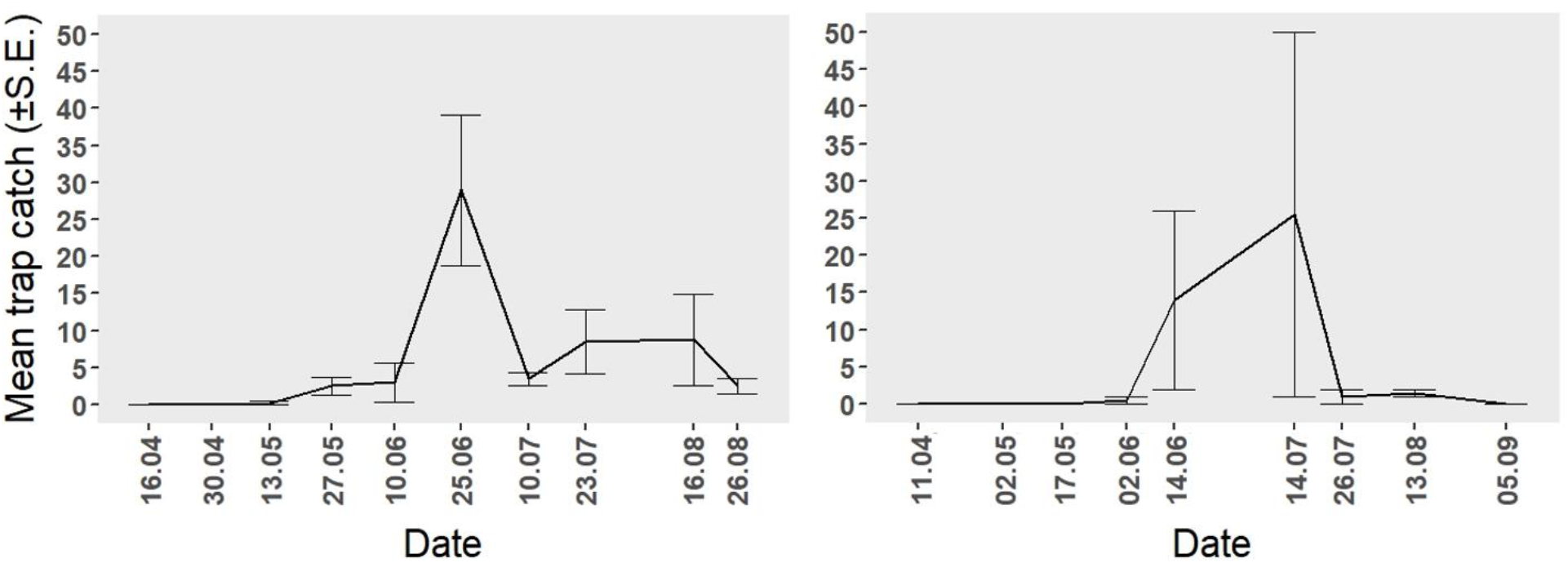
Mean trap captures of *Anisandrus maiche* in central-upper Ticino (left, Riviera-Iragna and Serravalle-Leggiuna, N = 4 ethanol-baited bottle traps) and in central Ticino (right, Bolette, Locarno, N=2 Polytraps).

### Other collection methods

No *A. maiche* were found infesting log sections (ca. 50 cm long, baited with ethanol) of European beech, Norway spruce and sweet chestnut at study sites (1)-(6) in Ticino. However, five specimens were collected from around and in the cork used to seal the ethanol reservoir of one log. No colonisation by *A. maiche* occurred of short sections of beech branches (about 20 cm long, 2-4 cm diameter, soaked in 70% ethanol) that had been placed on the ground at the same locations.

### *Phenology of* A. maiche *in Ticino*

Captures of *A. maiche* with ethanol-baited bottle traps in central-upper Ticino (Riviera-Iragna and Serravalle-Leggiuna) (n = 4 traps) revealed that the main flight period was from June (or late May) to August (Fig. 4). This general flight pattern was observed also with the captures with unbaited Polytraps which occurred from mid-May to mid-August (Fig. 4) near Locarno at locations that were mainly at slightly lower elevations than the bottle trap sites in central Ticino. As the Polytraps were used from 10 March (i.e., late winter/early spring) until 3 October (i.e., early autumn), this indicates that no flights were missed, and that there is no second flight period. Therefore, the species is probably univoltine.

### *Fungal associates of* A. maiche

From seven specimens of *A. maiche* that were caught alive, nine fungal cultures were successfully recovered. DNA barcoding confirmed that these belonged to the four following species: *Ambrosiella cleistominuta* (WSL DNA-ID PHP22_0914) (Ascomycota, Ceratocystidaceae; four cultures from sites Riviera-Iragna 2 and Serravalle-Leggiuna 1; **Fig. 5**), *Aureobasidium pullulans* (de Bary & Löwenthal) G. Arnaud (Ascomycota, Dothioraceae; three cultures), *Cladosporium cladosporioides* (Fresen.) G.A. de Vries (Ascomycota, Davidiellaceae; one culture), and *Fusarium lateritium* Nees (Ascomycota, Nectriaceae; one culture). *Cladosporium cladosporioides* was isolated from the beetle surface, whereas the other three species were isolated from crushed beetles.

**Figure 5.**
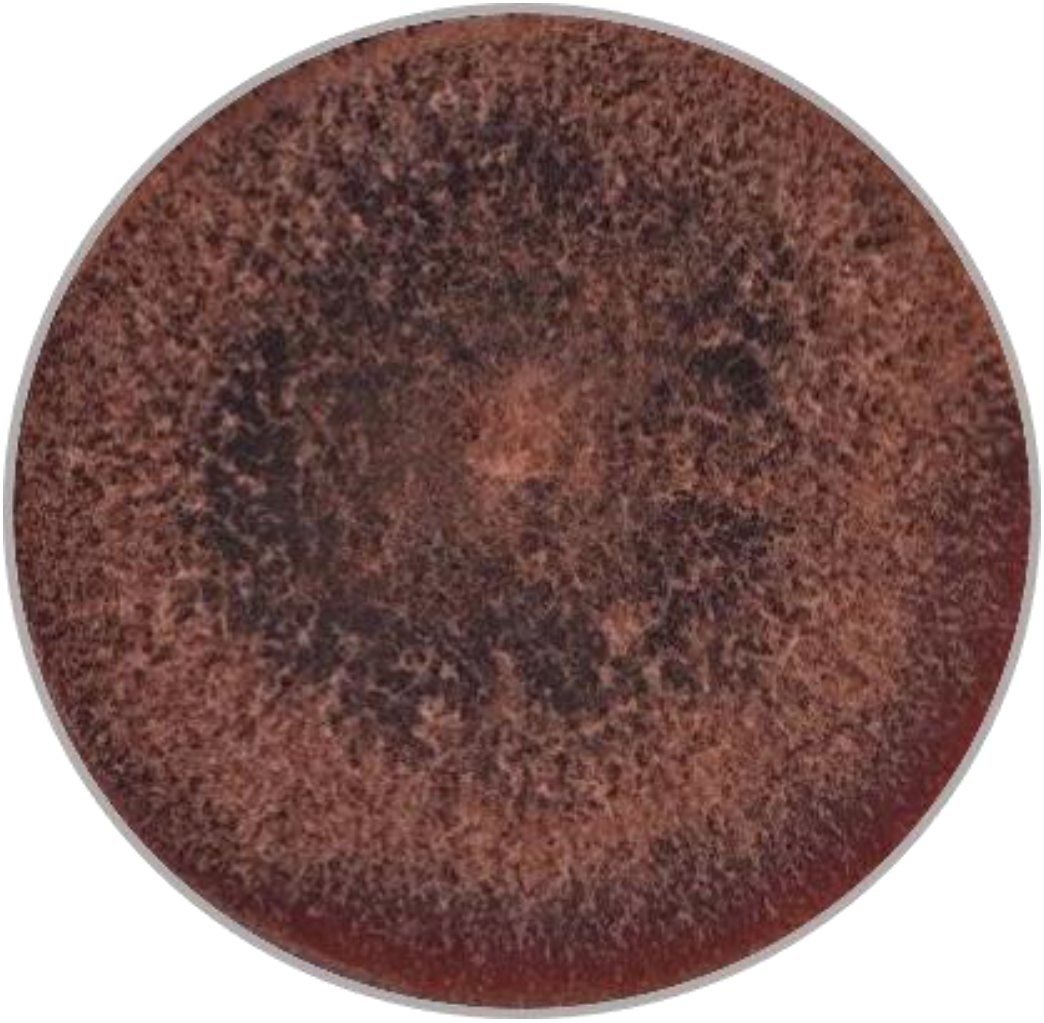
Culture of *Ambrosiella cleistominuta* from *Anisandrus maiche* trapped alive in Ticino.

### *Fungal associates of* X. crassiusculus

Cultures of fungal isolations from the mycangium of an adult *X. crassiusculus* (PHP22_1073– PHP22_1076) and from two breeding galleries (PHP22_0979, PHP22_1072) that were identified by DNA barcoding revealed that the associated fungus was *Ambrosiella cleistominuta*.

## Discussion

### *Establishment and distribution of* A. maiche *in Switzerland*

The detection of nearly 700 specimens of *Anisandrus maiche* across much of canton Ticino and an adjacent area of canton Grisons revealed that this species is established and already common in parts of southern Switzerland. This population appears to be contiguous with a recently detected population in adjacent parts of the Italian region of Lombardy where specimens of *A. maiche* were found about 30 km from the nearest known occurrence in Switzerland (Ruzzier et al. 2022). In northern Italy, *A. maiche* was found in five provinces between Milan and Treviso and appears to be relatively widespread even though the first detection of this species occurred only in 2021 (Colombari et al. 2022; Ruzzier et al. 2022). It is very likely that *A. maiche* has been present in northern Italy and southern Switzerland already for many years. Its resemblance to the common and widespread *Xylosandrus germanus* (Rabaglia et al. 2009), another non-native invasive ambrosia beetle of east Asian origin, probably prevented its earlier detection.

There is no indication that *A. maiche* occurs north of the Alpine divide (i.e., north of canton Ticino), although sampling and specific surveys for *A. maiche* north of the Alpine divide so far have occurred only in parts of cantons Valais and Zurich. However, given the abundance of *A. maiche* in Ticino and northern Italy and the considerable volume of international and domestic trade, it is probably only a matter of time until *A. maiche* is established north of the Alpine divide. Although its climatic requirements have not been determined thoroughly, the presence of *A. maiche* in Ukraine (Terekhova and Skrylnik 2012), western Russia near Moscow (Nikitskii 2009) and in far-eastern Russian Siberia (Kurentsov 1941) suggests that the climatic conditions north of the Alps should be suitable. However, species distribution modelling would be required to ascertain this with more certainty.

### *Phenology and host plants of* A. maiche

Captures of *A. maiche* in Ticino occurred between early May and late August 2022. Although this stretches across a period of nearly four months, there was no clear indication of two separate peaks of flight activity, and we assume that the species is univoltine. Based on observations in Ukraine, Skrylnik et al. (2019) also suggest that *A. maiche* is univoltine.

To our knowledge, no infestations by *A. maiche* of any trees have been reported from Ticino. Considering the wide distribution and apparent abundance in Ticino, this is surprising. However, based on host records from its native and non-native ranges (summarised in Hölling and Brockerhoff 2023 and Ruzzier et al. 2023), *A. maiche* is known to be highly polyphagous. In its native range in eastern Asia, host records include trees in the genera *Acer* (maple), *Alnus* (alder), *Betula* (birch), *Carpinus* (hornbeam), *Corylus* (hazel), *Fraxinus* (ash), *Juglans* (walnut), *Quercus* (oak), *Ulmus* (elm), and other genera including conifers (Hölling and Brockerhoff 2023). In the non-native range of *A. maiche* relatively few host records exist which, nonetheless, confirms that this species has a very wide host range. In Ukraine, attacks of *Betula* pendula, *Populus tremula, Quercus robur, Quercus rubra* (a North American oak species) and *Ulmus minor* have been reported (Nikulina et al. 2015; Skrylnik et al. 2019). In North America, *A. maiche* has been observed breeding in *Cercis canadensis, Cornus florida, Gleditsia triacanthos* and *Styrax japonicus* (Mayers et al. 2017; Ranger et al. 2015, 2019, 2020).

### *Introduction pathways for A. maiche* and other ambrosia beetles

To our knowledge, no confirmed border interceptions of *A. maiche* with any traded goods have been recorded, not in Switzerland, anywhere else within the EPPO regions nor in a number of other countries (Brockerhoff et al. 2006; Turner et al. 2021). However, many other ambrosia beetles including species in the genera *Xyleborus, Xyleborinus, Xylosandrus* and others have been intercepted numerous times (Brockerhoff et al. 2006; Haack et al. 2006, 2013), indicating the existence of pathways that can facilitate their invasions. For example, *Xylosandrus crassiusculus* and *Xylosandrus germanus*, two well-known invasive ambrosia beetles that are now established in several continents where they are not native, have been intercepted repeatedly in New Zealand (Brockerhoff et al. 2006). Furthermore, it is possible that specimens of *A. maiche* were intercepted but not recognised as this species (just as it was apparently overlooked in northern Italy and southern Switzerland until recently). Pathways known or thought to be involved in invasions of ambrosia beetles include international transport of wood packaging materials (such as pallets and case wood used with ceramic tiles, stone products and numerous other commodities) as well as trade in firewood and live plants (Liebhold et al. 2012; Meurisse et al. 2019). According to Mandelshtam et al. (2018), it may be possible that *A. maiche* extended its distribution by natural spread westwards from its native range in the Russian Far East; however, considering its discontinuous distribution between eastern Asia and central Europe, we regard it as more likely that international trade of infested wood or wood products is responsible for its arrival and establishment in Europe.

### Xylosandrus crassiusculus *in Switzerland*

The first detection in Switzerland of *X. crassiusculus*, an eastern Asian species, occurred in 2013 in southern Ticino where two specimens were found (Sanchez et al. 2020). We trapped 30 more specimens in 2022 at several locations between southern and central-upper Ticino (unpublished data), indicating that *X. crassiusculus* is also well established and relatively widespread. However, it appears to be considerably less abundant than *A. maiche*, at least at our trap locations. Even though the characteristic “frass noodles” produced by *X. crassiusculus* are highly visible (EFSA et al. 2020), the infestation of a *Hakea* shrub on the Brissago Island in Lago Maggiore is the first reported attack of a plant in Switzerland.

### *Fungal associates of* A. maiche *and* X. crassiusculus

*Ambrosiella cleistominuta* is the ambrosia fungus associated with *Anisandrus maiche* (Mayers et al. 2017). Previously, this association was only known from the United States, where the fungus was isolated and described as a species new to science in 2017 (Mayers et al. 2017). Here, we report for the first time that the fungus is also present in Europe, again in association with *A. maiche* (with four confirmed isolations from *A. maiche* in the present study). The fungus was most likely introduced to Europe together with its beetle mutualist. We are not aware of any studies indicating that this fungus is pathogenic to (live) plants.

Finding *A. cleistominuta* also in association with *Xylosandrus crassiusculus* in the samples of a *Hakea* shrub from the Brissago Island was unexpected. *Xylosandrus crassiusculus* is usually always associated with *Ambrosiella roeperi* (Harrington et al. 2014; Saragih et al. 2021). Ambrosia beetles and their mutualistic ambrosia fungi typically show a high degree of fidelity, and associations with other ambrosia fungi are rare (Biedermann and Vega 2020). However, new associations between ambrosia beetles and ambrosia fungi can occur and may be a result of invasions of ambrosia beetles into non-native regions (Menocal et al. 2023). Such new ambrosia symbioses may lead to reductions or even increases in beetle fitness (e.g., reproductive success) (Menocal et al. 2023). It is not known what the implications for *X. crassiusculus* of this novel symbiosis are; however, 10 adults of *X. crassiusculus* were found in the *Hakea* samples we obtained, which were able to complete their development.

Three other fungi were isolated from *A. maiche* trapped alive in Ticino: *Aureobasidium pullulans* is a saprophytic yeast-like fungus with a worldwide distribution; it occurs on the leaves of a wide range of plants and is known mainly from crops (Deshpande et al. 1992). *Cladosporium cladosporioides* is a member of a genus that includes the most common environmental fungi found worldwide with species of various lifestyles (Bensch et al. 2012). *Fusarium lateritium* is a globally distributed plant pathogen that may cause a variety of symptoms on affected plants. For example, *F. lateritium* can cause chlorotic leaf distortion on sweet potato (Clark et al. 1995) and nut grey necrosis on hazelnut fruits (Vitale et al. 2011). Recently, *F. lateritium* was found to cause shoot dieback of boxelder maple (*Acer negundo*) in Poland (Patejuk et al. 2022). It is not known whether this pathogen and its potential association with *A. maiche* could be a concern in Switzerland.

### *Potential damage to infested trees from* A. maiche *and potential tree pathogens*

So far, no trees or wood infested by *A. maiche* have been found in Switzerland, and it is uncertain what tree or shrub species are attacked in Switzerland. Based on previous host records and on the tree composition of sites where *A. maiche* was trapped in Ticino, this species may attack a wide range of broadleaved tree species present there. Most ambrosia beetles do not cause any damage of note, but there are exceptions, particularly when non-native ambrosia beetles and associated pathogenic fungi are involved (Eskalen et al. 2013; Hughes et al. 2017; Paap et al. 2018; Morales-Rodríguez et al. 2021). Based on observations from Ukraine, Terekhova & Skrylnik (2012) state that *A. maiche* “has no significant economic impact” in that country. However, “noticeable” damage to birch (*Betula pendula*) trees in Ukraine was described by Skrylnik et al. (2019) who gave *A. maiche* an intermediate impact rating for birch with a “physiological harmfulness score” of 5 out of 14, considering both damage from galleries made by the beetles and their assumed ability to act as vectors of plant pathogens. As of March 2023, there is no indication of any damage caused by *A. maiche* in Switzerland, neither from its galleries nor from any associated plant pathogens. However, no systematic surveys have been carried out yet, and it is not known if *A. maiche* is harmful to trees or shrubs. Nevertheless, its considerable abundance in some locations in Ticino along with the apparent association with at least on plant pathogen (*F. lateritium*, see Patejuk et al. (2022)) suggests that there is some potential for damage, especially if plants are stressed by drought, flooding or fire (Terekhova & Skrylnik 2012, Ranger et al. 2015, Mandelshtam et al. 2018). Ranger et al. (2015) consider that *A. maiche* prefers to attacks living, weakened trees over dead trees.

### Further research needs

There are several aspects of the invasion of *A. maiche* in Europe that deserve further study. As this species is already relatively common in southern Switzerland (and probably also in other invaded areas), it is possible that it will become one of the most abundant ambrosia beetles. This has happened at some locations in North America, where *A. maiche* was found to be one of the two most abundant ambrosia beetles, along with *Xylosandrus germanus* (Ranger et al. 2019). An increase in the abundance of *A. maiche* could lead to more noticeable damage of infested trees and it could also cause competition with native ambrosia beetles which may decline in response. Therefore, having a better understanding of the extent of the distribution of *A. maiche* beyond Ticino as well as knowledge of its host plants, fungal associates and their combined potential effect on plant health are desirable. Furthermore, a better understanding of the pathways involved with the invasions and spread of *A. maiche* and other ambrosia beetles could be used to limit the extent of future invasions and impacts. Finally, given the occurrence of heteroplasmy in COI sequences in some of the samples of *A. maiche* we analysed, a known problem with COI barcoding of insects (e.g., Magnacca and Brown 2010; Cognato et al. 2020), further studies on the invasion and population genetics of *A. maiche* should examine the use of additional markers.

## Abbreviations used

FPS: Forest Protection Switzerland (Waldschutz Schweiz), Swiss Federal Institute for Forest, Snow and Landscape Research WSL, Birmensdorf, Switzerland
MSNL: Museo di storia naturale (Natural History Museum), Lugano, Switzerland
PHP: Phytopathology, Swiss Federal Institute for Forest, Snow and Landscape Research WSL, Birmensdorf, Switzerland
UPN: Ufficio della Natura e del Paesaggio, Bellinzona, Switzerland
WSL: Swiss Federal Institute for Forest, Snow and Landscape Research WSL, Birmensdorf, Switzerland

## Acknowledgements

We thank Adrian Oncelli (Repubblica e Cantone Ticino, Dipartimento del territorio, Sezione forestale) for logistical support and contributions to trapping, Aline Knoblauch (Office fédéral de l’environnement (OFEV), Division Forêts) for project support, Vienna Kowallik (Universität Freiburg, Forstentomologie und Waldschutz) for training in fungal extraction, Quirin Kupper, Sven Ulrich und Robin Winiger (WSL) for laboratory support, Carolina Cornejo (WSL) for interpretation of DNA sequences, and Carl-Michael Anderson for taking photos of *A. maiche*. This project was supported in part by funding from the Swiss Federal Office for the Environment (FOEN/OFEV/BAFU) to WSL. MK was partly supported by the Ministry of Agriculture of the Czech Republic, institutional support MZE-RO0118.

## Notes

### Competing Interest Statement

The authors have declared no competing interest.

